# Defective mesenchymal Bmpr1a-mediated BMP signaling causes congenital pulmonary cysts

**DOI:** 10.1101/2023.09.26.559527

**Authors:** Yongfeng Luo, Ke Cao, Joanne Chiu, Hui Chen, Hong-Jun Wang, Matthew E. Thornton, Brendan H. Grubbs, Martin Kolb, Michael S. Parmacek, Yuji Mishina, Wei Shi

**Affiliations:** Department of Surgery, Children’s Hospital Los Angeles, Keck School of Medicine, University of Southern California, Los Angeles, CA 90027; Division of Pulmonary, Critical Care & Sleep Medicine, Department of Internal Medicine, University of Cincinnati College of Medicine, Cincinnati, OH 45267, USA; Division of Maternal Fetal Medicine, Department of Obstetrics and Gynecology, Keck School of Medicine, University of Southern California, Los Angeles, CA 90089, USA; Department of Medicine, McMaster University, Hamilton, ON, Canada L8N 4A6; Department of Medicine, Perelman School of Medicine, University of Pennsylvania, Philadelphia, PA 19104, USA; Department of Biologic and Material Sciences, University of Michigan, 1011 N. University Ave., Ann Arbor, MI 48109

**Keywords:** BMP signaling, Bmpr1a, Lung mesenchymal cells, Airway smooth muscle cells, Pulmonary cysts, Lung development, Lung branching morphogenesis

## Abstract

Abnormal lung development can cause congenital pulmonary cysts, the mechanisms of which remain largely unknown. Although the cystic lesions are believed to result directly from disrupted airway epithelial cell growth, the extent to which developmental defects in lung mesenchymal cells contribute to abnormal airway epithelial cell growth and subsequent cystic lesions has not been thoroughly examined. In the present study, we dissected the roles of BMP receptor 1a (Bmpr1a)- mediated BMP signaling in lung mesenchyme during prenatal lung development and discovered that abrogation of mesenchymal *Bmpr1a* disrupted normal lung branching morphogenesis, leading to the formation of prenatal pulmonary cystic lesions. Severe deficiency of airway smooth muscle cells and subepithelial elastin fibers were found in the cystic airways of the mesenchymal *Bmpr1a* knockout lungs. In addition, ectopic mesenchymal expression of BMP ligands and airway epithelial perturbation of the Sox2-Sox9 proximal-distal axis were detected in the mesenchymal *Bmpr1a* knockout lungs. However, deletion of Smad1/5, two major BMP signaling downstream effectors, from the lung mesenchyme did not phenocopy the cystic abnormalities observed in the mesenchymal *Bmpr1a* knockout lungs, suggesting that a Smad-independent mechanism contributes to prenatal pulmonary cystic lesions. These findings reveal for the first time the role of mesenchymal BMP signaling in lung development and a potential pathogenic mechanism underlying congenital pulmonary cysts.

## Introduction

Congenital pulmonary cysts, resulting from abnormal fetal lung development, causes respiratory distress, infection, and pneumothorax in neonates. The dynamic pathogenic process and mechanisms are difficult to study in humans and few animal models are available. It is known that lung development begins with the specification of the respiratory domain in the ventral wall of the anterior foregut endoderm, as indicated by the expression of Nkx2-1, and the separation of the respiratory tract from the dorsal esophagus. The primary lung epithelial buds then undergo reiterated elongation and division to form the conducting airways (branching morphogenesis), followed by distal saccular formation and peripheral alveolarization to give rise to millions of gas exchange units ^1,2^. This well-known process literally describes the growth of respiratory epithelium based on the extensive studies done in the past decades. It is notable that lung morphogenesis is a complex process relying on the highly coordinated development of both lung epithelium and lung mesenchyme. Evidence has increasingly suggested that lung mesenchymal lineages, including airway and vascular smooth muscle cells, pericytes, and stromal fibroblasts, are as important as epithelial cells in proper lung development ^3-5^. In addition to acting as a mechanical framework that supports the formation of the bronchi, bronchioles, and the distal alveoli, the lung mesenchyme provides a microenvironment that regulates the growth of epithelium through a variety of morphogenic signals, such as Wnts, fibroblast growth factors (Fgfs), and bone morphogenetic proteins (BMPs) ^1,6-8^.

BMPs are a family of growth factors that direct many biological processes, including organogenesis ^9^. BMPs bind to the receptor complex of BMP receptor II (Bmpr2) and BMP receptor I (Bmpr1a or Bmpr1b), which in turn activates the intracellular Smad-dependent and Smad-independent pathways ^10^. Mice with conventional *Bmpr1a* gene deletions are early embryonic lethal (E7.5-9.5) before lung organogenesis ^11^. As reported by us and other groups, both hyperactivation and inhibition of BMP signaling in fetal lung epithelial cells result in lung malformation, which is primarily due to the defects in distal lung epithelial cell proliferation and differentiation/maturation ^12-15^. However, the role of mesenchymal BMP signaling in regulating fetal lung development has not been studied due to the lack of lung mesenchyme-specific targeting tools in the past. We recently generated a lung mesenchyme-specific *Tbx4* lung enhancer-driven *loxP*/Cre mouse driver line ^16^ which enables us to manipulate BMP signaling specifically in lung mesenchymal cells in vivo and to study its role in the lung development.

Although Bmpr1a is expressed predominantly in fetal mouse lung airway epithelial cells at the early gestation stage (embryonic day (E)12.5 to E14.5), its expression in fetal lung mesenchyme is also evident during mid gestation ^13^. Herein, we specifically deleted *Bmpr1a* in fetal lung mesenchymal cells, which resulted in abnormal airway development and subsequent prenatal cystic malformation. This pathological phenotype resembles the features observed in pediatric patients diagnosed with congenital pulmonary airway malformation (CPAM). Therefore, investigating the abnormal lung phenotypes caused by lung mesenchyme-specific *Bmpr1a* knockout and revealing the underlying molecular and cellular mechanisms will significantly enhance our understanding of congenital lung diseases.

## Results

### Abrogation of Bmpr1a-mediated BMP signaling in lung mesenchyme disrupted fetal lung development

By crossing the *Tbx4-rtTA/TetO-Cre* driver line to the *floxed-Bmpr1a* mice (*Tbx4-rtTA/TetO-Cre/Bmpr1a^fx/fx^*), *Bmpr1a* was specifically knocked out in the lung mesenchyme with doxycycline (Dox) induction from the beginning of lung formation (E6.5, Suppl. Fig.1A). This was validated at the mRNA and protein levels (Suppl. Fig.1B and 1C). For example, in E15.5 mesenchyme-specific *Bmpr1a* conditional knockout (CKO), Bmpr1a immunostaining was absent in lung mesenchyme but present in the epithelia (Suppl. Fig.1C) while Bmpr1a was detected in both mesenchyme and epithelia of wildtype (WT) lungs. By gross view with quantitative analysis (Fig.1A-C), early embryonic lung morphogenesis between *Bmpr1a* CKO lungs and WT controls was comparable prior to E13.5. Decreased epithelial branching and increased size of branching tips were observed from E14.5. As lung development progressed, the terminal airways in *Bmpr1a* CKO lungs displayed dilation and exhibited increasingly extensive cystic changes. However, the overall size of the lungs remained relatively unchanged. The dynamic change of the lung cysts in *Bmpr1a* CKO lungs was further analyzed in H&E-stained tissue sections (Fig.1D). The walls of the lung cysts were lined with a single layer of epithelial cells, over the course of development, ultimately resulting in the formation of balloon-like cystic lesions by the end of gestation (E18.5). These findings were consistent with the overall morphology of the lungs. Interestingly, overall cell proliferation and apoptosis at E15.5, when lung cysts were developed, had no significant change between cystic *Bmpr1a* CKO and WT control lungs as measured by EdU labeling and TUNEL assay, respectively (Fig.1E-G).

**Fig. 1.**
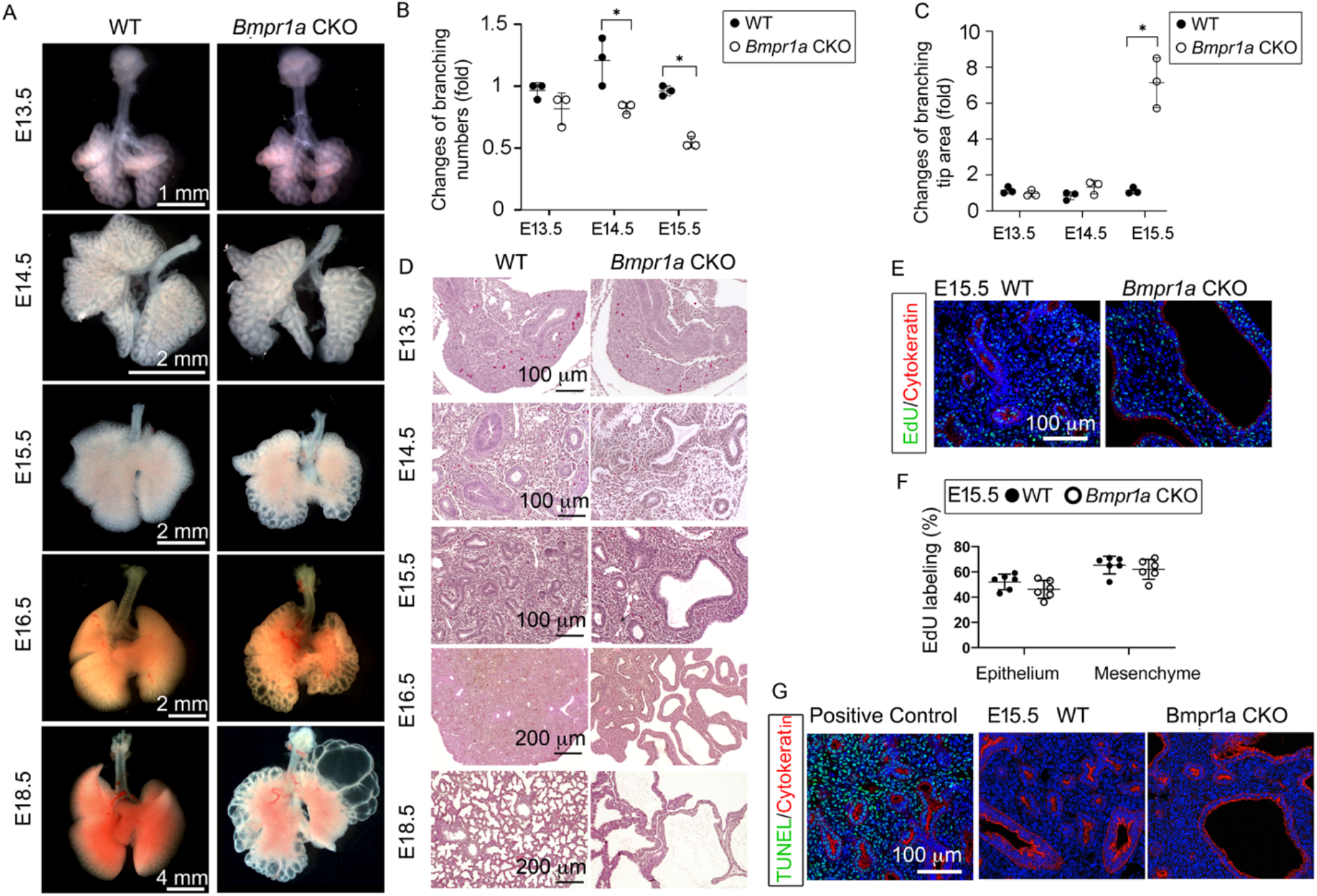
Lung mesenchyme-specific deletion of *Bmpr1a* caused abnormal lung morphogenesis and prenatal airway cystic lesions beginning in mid-gestation. (A) Brightfield images of whole WT and *Bmpr1a* CKO mouse lungs at different embryonic stages. (B & C) Quantitative measurement and comparison of terminal airway branching numbers and sizes. (D) H&E-stained *Bmpr1a* CKO lungs at different embryonic stages. (E & F) EdU incorporation study for cell proliferation analysis in lung mesenchymal and cytokeratin-positive epithelial cells (n=6). (G) Apoptosis analysis by TUNEL assay. The positive control slides for apoptosis were generated by treating the tissue sections with DNase I. Pictures are representative of at least five samples in each condition.

### Deletion of Bmpr1a in fetal lung mesenchymal cells resulted in deficiency of airway-specific smooth muscle growth

To understand the molecular mechanisms underlying the aforementioned phenotypic changes, bulk RNA-seq was used to examine the differentially expressed genes (DEGs) between *Bmpr1a* CKO and WT lung tissues at E15.5. A total of 1,001 DEGs (FDR ≤ 0.05 and log_2_FC ≥ 1) were identified, among which 547 genes were upregulated and 454 genes were downregulated in *Bmpr1a* CKO lungs as compared to the transcriptome of WT lungs (Fig.2A). The raw data are available in the Gene Expression Omnibus repository under accession number GSE97946. Gene Ontology (GO) enrichment analysis showed that lung mesenchymal *Bmpr1a* regulates a large number of genes involved in the Muscle System Process (*P* < 0.001, Fig. 2B).

**Fig. 2.**
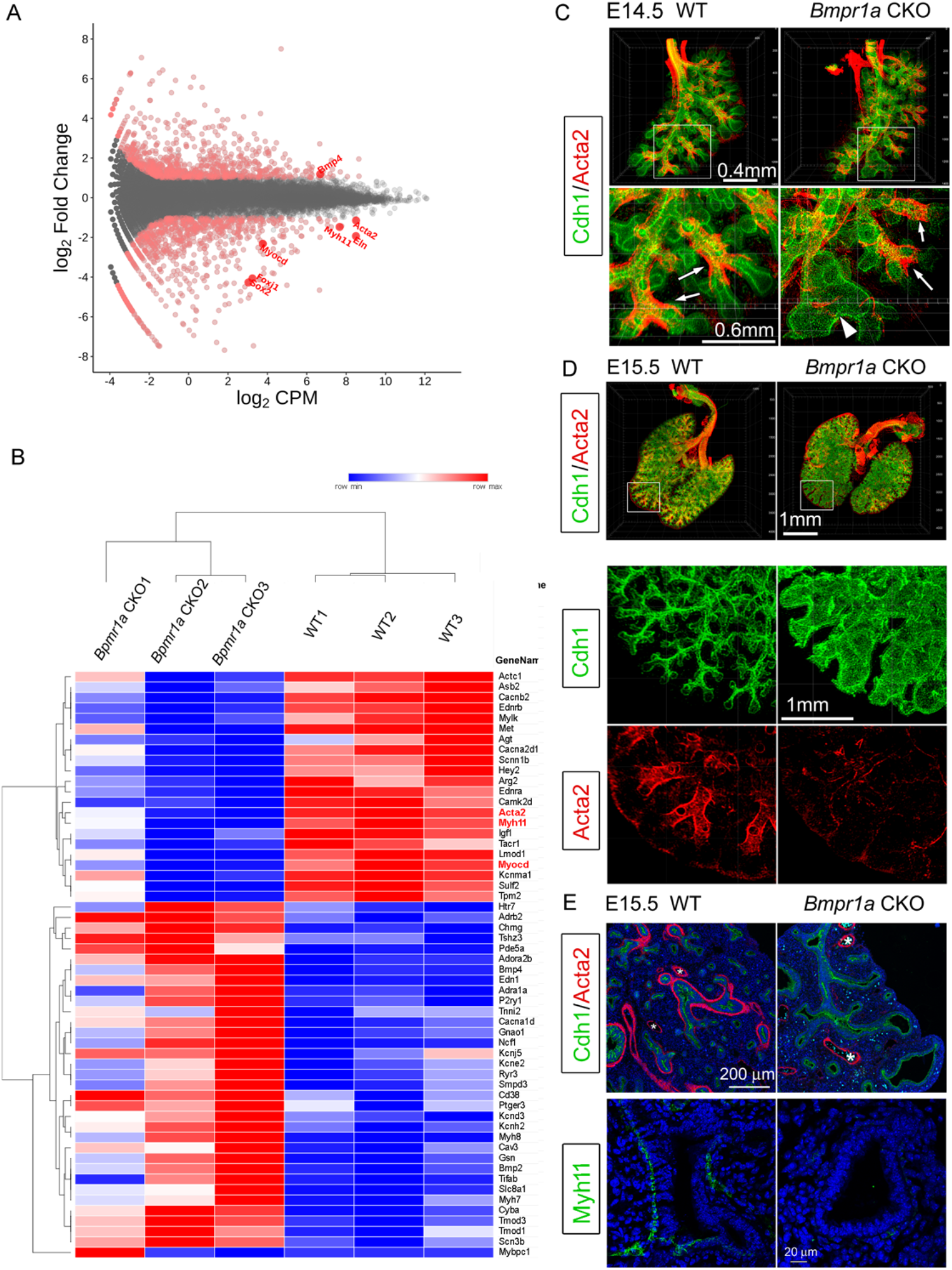
Airway smooth muscle was substantially reduced in *Bmpr1a* CKO lungs. (A) Scatter plots of RNA-Seq analysis for DEGs (n=3). (B) RPKM heatmap of Muscle System Process Genes (GO:0003012). Genes involved in the Muscle System Process with significant changes between *Bmpr1a* CKO and WT lungs are shown in the RNA-seq heatmap. The input data was RPKM values generated from the raw data using the R/Bioconductor program ‘edgeR’. (C & D) Whole mount immunostaining of Cdh1 and Acta2 in E14.5 and E15.5 lungs. Excessive dilation of airway terminals and extensive reduction of airway smooth muscle were observed in E15.5 *Bmpr1a* CKO lungs. The normal-looking airways are marked by arrows and the enlarged epithelial bud accompanied with compromised smooth muscle are marked by arrowheads. (E) E15.5 lung section stained with Cdh1& Acta2 or Myh11. *Vascular structures.

As shown above in Fig.1, enlarged airways were initially seen in *Bmpr1a* CKO lungs from E14.5. Whole mount staining of smooth muscle actin and comprehensive 3D imaging revealed a specific absence of airway smooth muscle cells (SMCs) surrounding the dilated airway branching in *Bmpr1a* CKO lungs, while unaffected airway branches in *Bmpr1a* CKO lungs remained surrounded by SMCs, as compared to the WT lungs (Fig. 2C). As lung development proceeded to E15.5, the distal airway enlargement became increasingly severe and extensive, as indicated by the whole mount staining of Cdh1 (E-Cadherin), which outlines the airway epithelia. Reduction and even lack of airway SMCs was consistently found around the enlarged airways of E15.5 *Bmpr1a* CKO lungs, shown by immunostaining for smooth muscle cell markers Acta2 and Myh11 (Fig. 2D). In contrast, vascular SMCs in *Bmpr1a* CKO lungs appeared to remain unaffected (Fig. 2E).

### The effect of Bmpr1a deletion on the other components of lung mesenchyme

The *Tbx4* driver line also targets lung pericytes and endothelial cells with early induction ^16^. However, *Bmpr1a* deletion in these cell lineages had no effects on these cells or the associated pulmonary vasculature, as measured by the relevant cellular markers (Cspg4 and Pecam1) at both the mRNA and protein levels (Fig.3A and B). Emerging evidence has suggested that the extracellular matrix (ECM), including the airway basement membrane and matrix fibril collagen, is critical to lung development and maturation ^8,17^. Several major components of the ECM in the *Bmpr1a* CKO lungs, such as laminin and type III collagen, were examined at both the mRNA and protein levels. No significant changes were found, even in the areas surrounding the dilated airways and lung cysts when compared to the WT lungs (Fig.3A-3B).

**Fig. 3.**
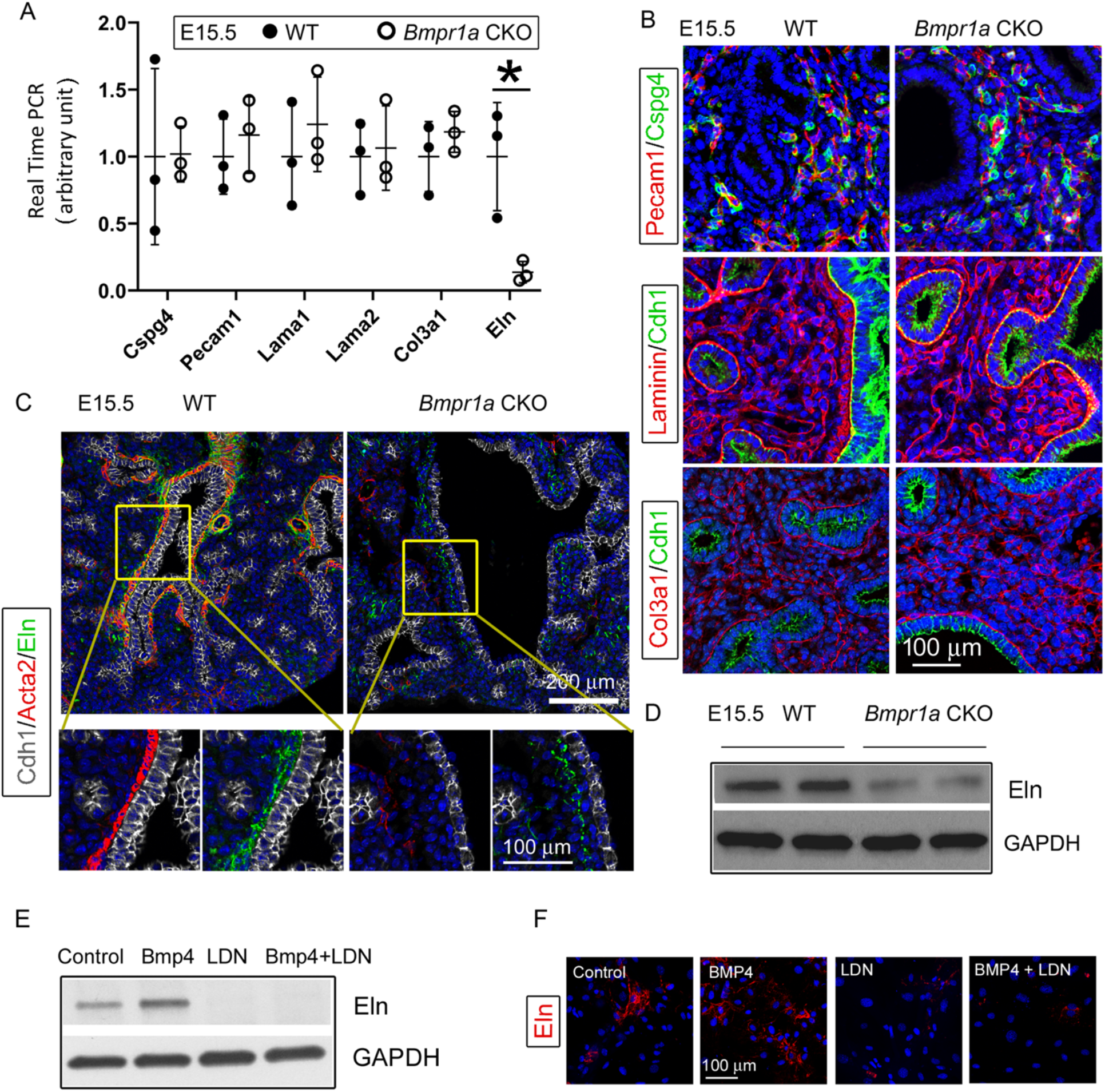
Bmpr1a CKO resulted in a significant reduction in elastin expression underneath the airway epithelia of E15.5 lungs, but not in the pericytes, vasculature, and basement membrane. (A) Expression of *Cspg4*, *Pecam1*, *Lama1*, *Lama2*, *Col3a1* and *Eln* at the mRNA level was measured by real-time PCR, \**P* < 0.05. (B) E15.5 lung section stained with Cspg4, Pecam1, Laminin and Col3a1. (C) E15.5 lung section stained with Cdh1, Acta2, and Elastin. (D) Reduced elastin expression of *Bmpr1a* CKO lungs at the protein level was detected by WB. (E-F): Elastin expression in E15.5 lung mesenchymal cells was upregulated by BMP4 and downregulated by BMP type 1 receptor-specific inhibitor LDN193189 (LDN), as detected by WB and immunofluorescence staining.

However, the mesenchymal knockout of *Bmpr1a* caused significant reduction of elastin expression, as detected by real-time PCR and western blot (Fig3.A and D). In the *Bmpr1a* CKO lungs, co-immunostaining of epithelia (Cdh1), smooth muscle cells (Acta2), and elastin (Eln) showed that this decrease in elastin expression occurred primarily in the areas adjacent to the epithelia of enlarged airways with the deficiency of airway smooth muscle. This suggests that the elastin abnormality may be caused by airway SMC deficiency or by reduced elastin expression in adjacent epithelial cells (Fig.3C). To determine whether BMP signaling can directly regulate elastin expression in fetal lung mesenchymal cells, alteration of elastin protein expression upon BMP4 treatment in primary fetal lung mesenchymal cells was analyzed. A significant increase in elastin protein was observed in the cells treated with BMP4 (Fig.3E-3F). However, inclusion of the BMP type 1 receptor (BMPR1) inhibitor LDN193189 (LDN) fully blocked the BMP4-stimulated elastin upregulation.

### Bmpr1a-mediated BMP signaling regulated airway SMC differentiation through a Smad independent pathway

Activation of the BMP receptor complex can trigger intracellular Smad-dependent and Smad-independent pathways ^10^. In line with this report, the downregulation of Smad1/5-mediated BMP signaling was also observed in the *Bmpr1a* CKO lungs, evident from the reduced phosphorylation of Smad1/5 (p-Samd1/5 in Fig.4A). The role of Bmpr1a in mediating Smad1/5 activation in lung mesenchymal cells was further investigated in primary culture of fetal lung mesenchymal cells isolated from WT fetal lung tissues. Treatment of these cell with BMP4 activated Smad1/5 phosphorylation, while the addition of BMPR1 inhibitor LDN reduced the BMP4-induced Smad1/5 phosphorylation (Fig. 4E). To determine whether the Smad1/5-mediated BMP canonical pathway controls airway SMC growth in vivo, *Smad1* and *Smad5* were both deleted in lung mesenchyme by crossing the floxed-*Smad1* and -*Smad5* mice with the *Tbx4-rtTA/TetO-Cre* driver. Although abnormal fetal lungs with hypoplastic development were present in the *Smad1/5* double CKO mice, airway dilation and cystic lesions were not detected (Fig. 4B-4C). Furthermore, in the *Smad1*/*5* CKO lungs, airway SMC growth was not affected, as indicated by the expression of Acta2 around the airways (Fig.4D). This different phenotypic observation suggests that airway SMC growth may be independent of Smad1/5-mediated pathway.

**Fig. 4.**
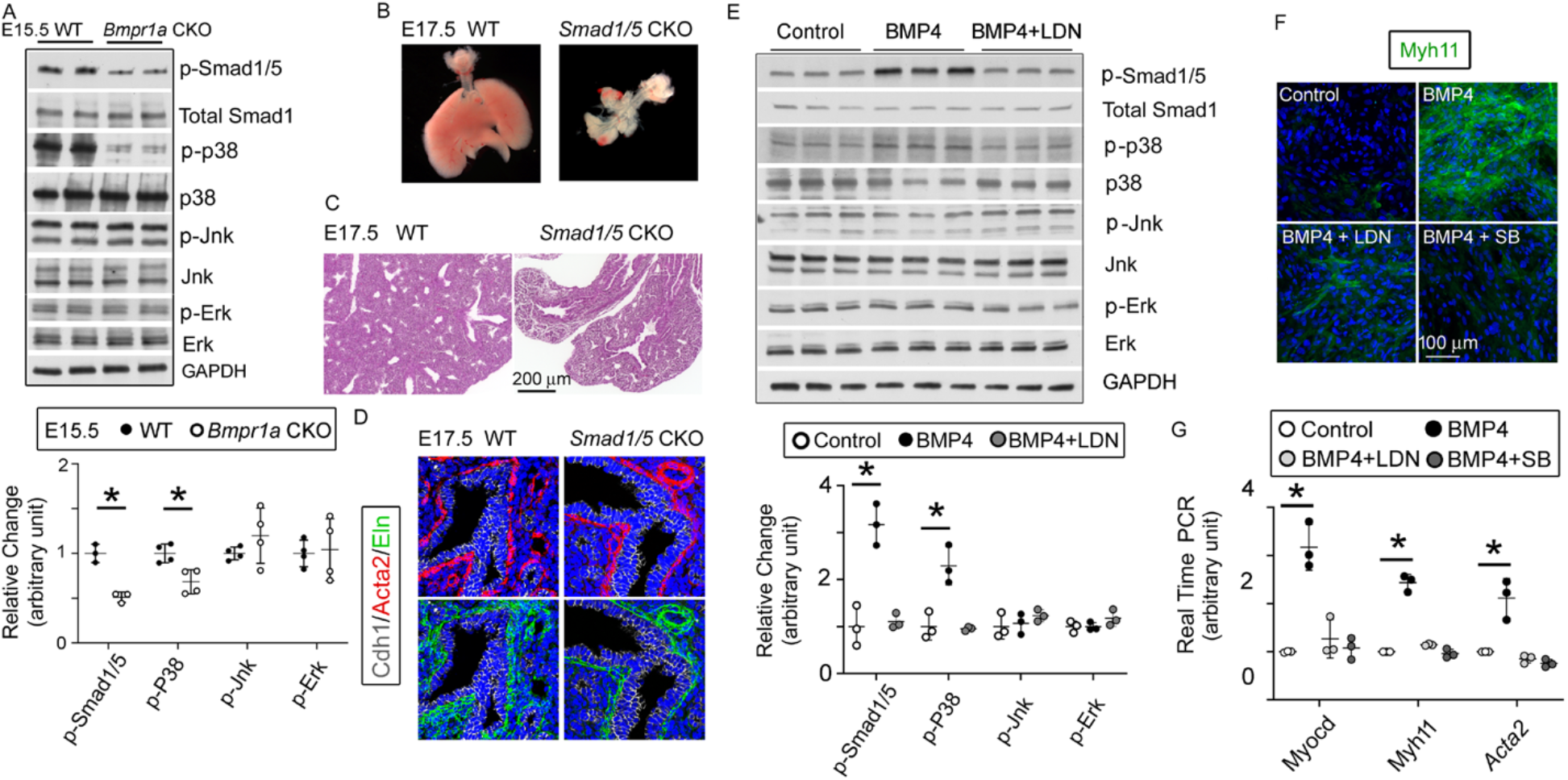
The BMP pathway regulates the myogenesis of lung mesenchymal cells via the Smad independent pathway. (A) Activation of the intracellular downstream Smad1, p38, Jnk and Erk signaling pathways in WT and *Bmpr1a* CKO lung tissues was detected by the WB and quantified by densitometry. The levels of protein phosphorylation were normalized by the corresponding total protein and is presented as a relative change to the WT, \**P* < 0.05. (B) Gross view of whole lungs from *Smad1*/*5* double conditional knockout mice (*Smad1/5* CKO) and WT littermates showed that simultaneous deletion of *Smad1* and *Smad5* in lung mesenchyme completely disrupted lung development. (C) No airway dilation or cysts were observed in the H&E-stained tissue sections of the *Smad1/5* CKO lungs at E17.5. (D) Expression of airway SMCs and elastin was not altered in E17.5 *Smad1/5* CKO lungs, as shown by immunostaining of Cdh1, Acta2, and elastin. (E) Changes of intracellular signaling pathways in cultured fetal lung mesenchymal cells upon treatment with BMP4 (50ng/ml) and/or LDN193189 (200 nM) was detected by WB and quantified by densitometry. The relative change to the control condition is presented, \**P* < 0.05. (F-G) Altered expression of SMC genes at the protein level (Myh11) and the mRNA level (*Myocd, Myh11 and Acta2*) was respectively analyzed by immunostaining and real-time PCR for the primary culture of E15.5 WT lung mesenchymal cells treated with BMP4 (50ng/ml), LDN (200 nM) and SB (1 µM), \**P* < 0.05.

We then examined other major Smad-independent pathways in the *Bmpr1a* CKO lungs. Reduced phosphorylation of p38 (P-p38) MAP Kinase (MAPK), but not Jnk and Erk, was detected in the *Bmpr1a* CKO lungs (Fig.4A). Bmp4 treatment of the cultured primary lung mesenchymal progenitor cells also specifically activated p38, a response that was specifically blocked by the addition of the BMPR1 inhibitor LDN (Fig.4E). Additionally, the role of the BMP4-Bmpr1a-p38 pathway in regulating SMC-related gene expression was assessed in cultured fetal lung mesenchymal cells. As previously reported ^18^, Myocd is a key transcription co-factor in controlling airway SMC differentiation. Myh11 and Acta2, major contractile proteins, are two prominent SMC markers. Treatment of lung mesenchymal progenitor cells with BMP4 induced a pronounced increase in the expression of these SMC-related genes at both the mRNA and protein levels (Fig.4F-4G). Inhibition of either Bmpr1a by LDN or p38 by SB203580 (SB) significantly attenuated the promoting effect of BMP4 on SMC differentiation.

### Bmpr1a-mediated Bmp signaling specifically promoted non-vascular smooth muscle cell growth

As shown in Fig. 2E, *Bmpr1a* CKO caused defective growth of airway SMCs in vivo, while vascular SMCs remained unaffected. To investigate the distinct effects of the Bmpr1a-mediated pathway on non-vascular SMCs vs. vascular SMCs, primary fetal lung mesenchymal cells were isolated from *Tagln-YFP*/*Cspg4-DsRed* double reporter mice, in which airway SMCs/myofibroblasts were solely labeled with YFP expression while vascular SMCs and pericytes were marked by both YFP and DsRed expression (Fig.5A). Following fluorescence-activated cell sorting (FACS) of the single cell suspension obtained from dissociated E15.5 lung tissues (Fig.5B), the vascular SMCs (YFP^+^/DsRed^+^) were separated from non-vascular SMCs (YFP^+^/DsRed^-^). These two distinct SMC populations were then cultured for further analysis ^19^. As shown in Fig. 5C, BMP4 treatment could promote SMC-associated gene expression in lung SMCs of non-vascular origin rather than those of vascular origin, as assessed by their Myh11 expression. This BMP4-induced upregulation of Myh11 expression in non-vascular SMCs was effectively blocked by concurrent treatment with the BMP type 1 receptor-specific inhibitor LDN.

**Fig. 5.**
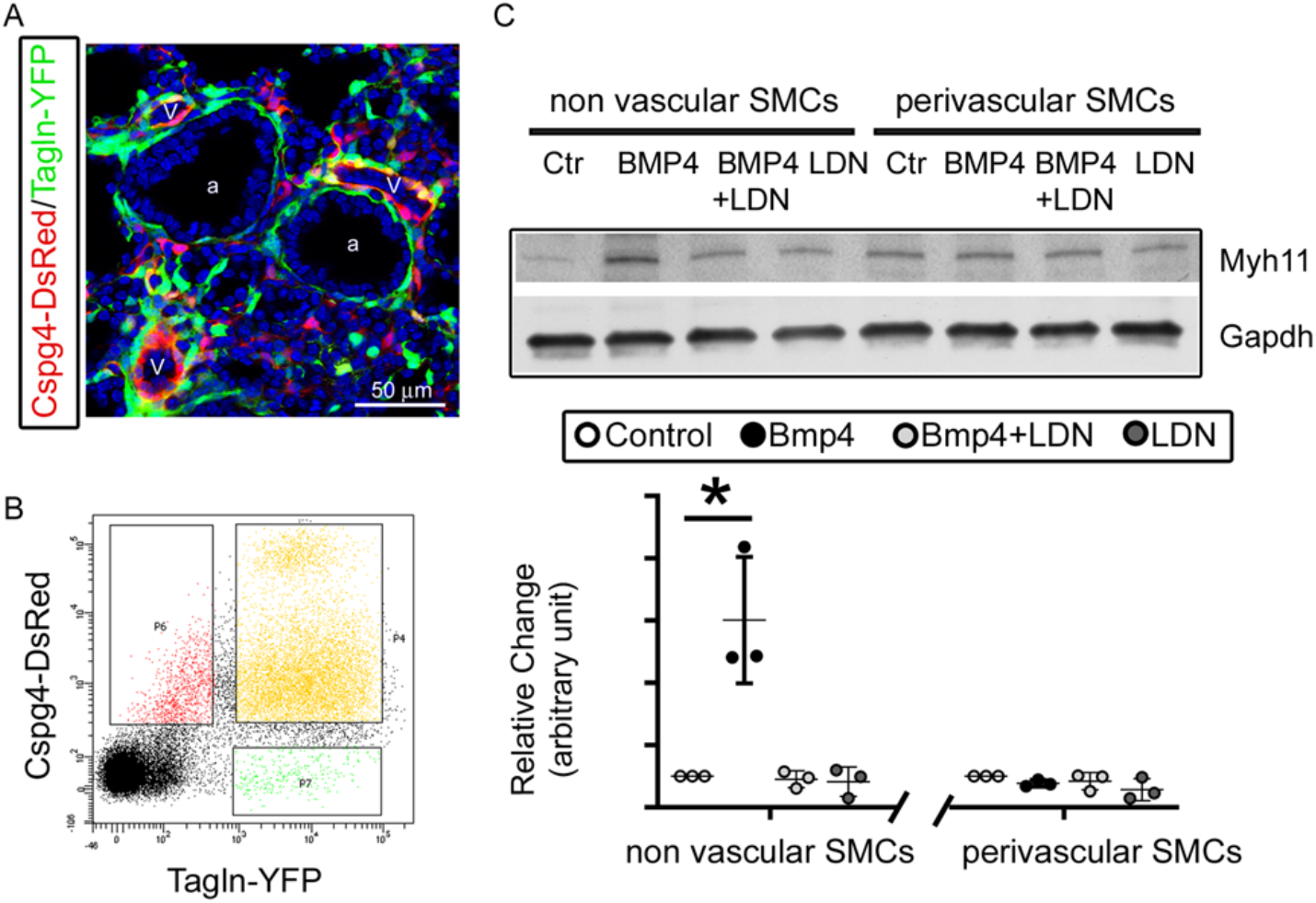
Bmpr1a-mediated signaling played different roles in the differentiation of airway versus vascular SMCs. (A) YFP and DsRed expression patterns in the fetal lungs of *Tagln-YFP*/*Cspg4-DsRed* mice. a: airway, v: vessel. (B) Airway SMCs (YFP^+^) and vascular SMCs (YFP^+^/DsRed^+^) were isolated by FACS sorting. (C) The differential effect of Bmp4 treatment (50 ng/ml) on contractile protein Myh11 expression was analyzed in SMCs of non-vascular origin (YFP^+^) versus vascular origin (YFP^+^/DsRed^+^), and the role of Bmpr1a in mediating this effect was tested by adding its specific inhibitor LDN193189 (200 nM). The WB data of Myh11 expression was normalized to GAPDH (loading control) and is represented as a relative change to the control condition, \**P* < 0.05.

### Loss of airway SMCs alone was not sufficient to cause prenatal cystic malformation in vivo

Previous studies using embryonic lung explant culture have suggested that airway SMCs play an important role in branching morphogenesis in developmental lungs ^20,21^. *Myocd* encodes an important transcriptional coactivator of serum response factor (SRF), modulating the expression of smooth muscle-specific cytoskeletal and contractile proteins ^22^. In our *Bmpr1a* CKO lungs, expression of *Myocd* was substantially downregulated (Fig.6A). To determine whether Bmpr1a-mediated downregulation of Myocd expression and the resultant airway SMC defects directly caused fetal airway cystic lesions in vivo, *Myocd* was deleted in lung mesenchyme starting at E6.5 using the same *Tbx4-rtTA/TetO-Cre* driver (Suppl. Fig.2). As detected by Acta2 immunostaining, knockout of *Myocd* led to a severe deficiency in airway SMC development (Fig.6B) resembling the airway SMC deficiency observed in the *Bmpr1a* CKO lungs (Fig. 2 and 3C). However, mesenchymal Bmpr1a expression remained unchanged in the *Myocd* CKO lungs. No significant branching defect or cystic malformation was observed in the *Myocd* CKO lungs (Fig.6C-6D), which is consistent with the previous report ^18^. Despite severe disruption of airway SMC development in the *Myocd* CKO lungs, elastin fiber deposition in the airways was unaffected (Fig. 6B), which was a significant difference when compared to the findings in the *Bmpr1a* CKO airways, where a deficiency of elastin fibers adjacent to the epithelium was observed (Fig. 3C).

**Fig. 6.**
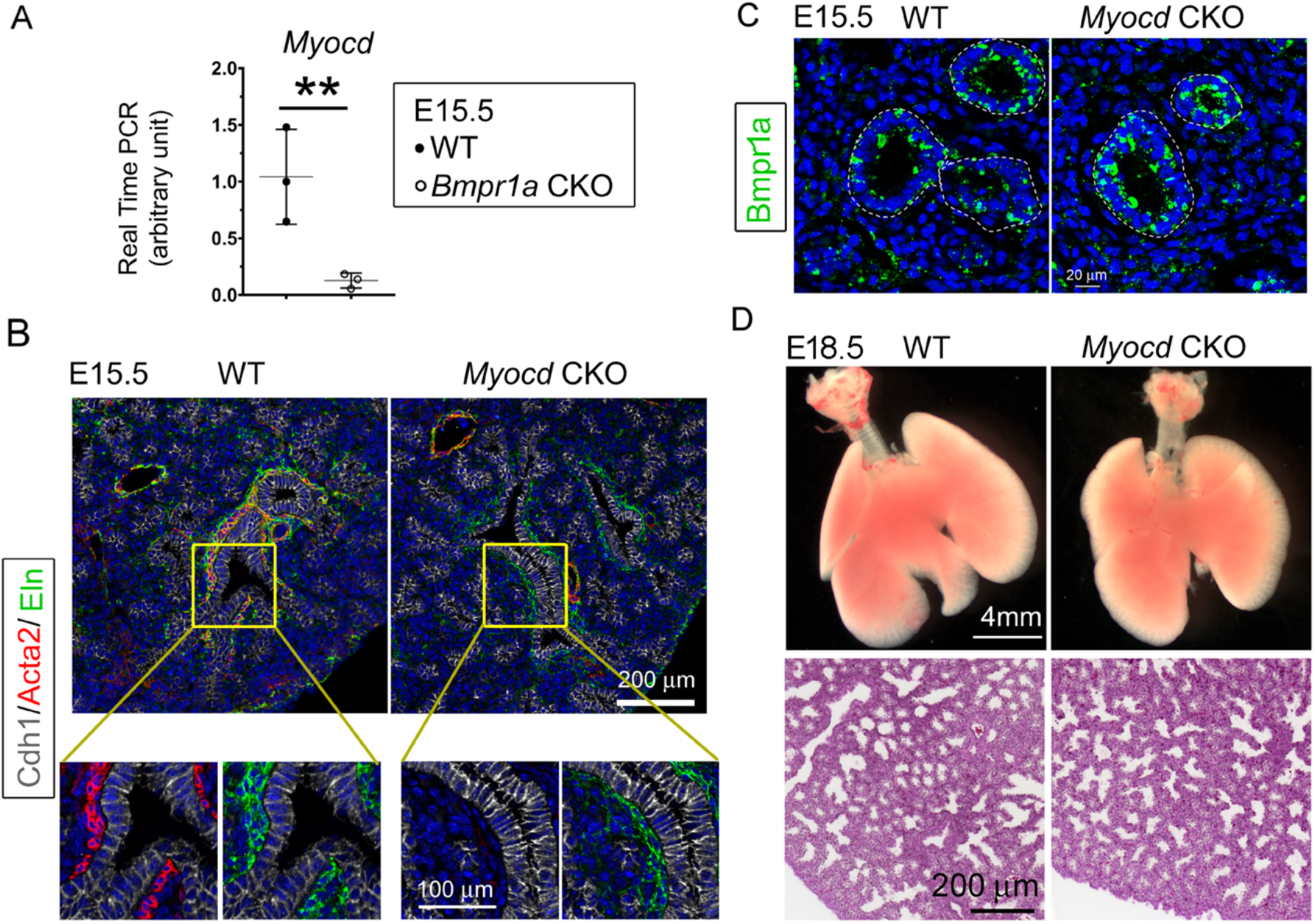
Lung mesenchymal knockout of *Myocd* did not cause any branching abnormalities or lung cysts. (A) The expression of *Myocd* in *Bmpr1a* CKO lungs was substantially decreased, as measured by real-time PCR, \*\**P* < 0.01. (B) Deficiency in airway SMCs was observed in E15.5 mesenchyme-specific *Myocd* CKO lungs, as shown by co-immunofluorescence staining of Cdh1, Acta2, and elastin. (C) Comparison of Bmpr1a expression between E15.5 *Myocd* CKO and WT control lungs by immunofluorescence staining. (D) Comparison of the lungs between WT and *Myocd* CKO mice at the end of gestation (E18.5) did not reveal any significant morphological changes by gross view. No histological difference was found between the WT and the *Myocd* CKO lungs by examining their H&E-stained lung tissue sections.

### The proximal-distal developmental patterning of fetal lung epithelia was perturbed in the cystic lungs of the *Bmpr1a* CKO mice

The coordinated airway epithelial differentiation that forms the proximal-distal axis is another important mechanism controlling lung morphogenesis ^23^. The proximal epithelial cells, marked by the expression of several unique genes such as Sox2, serve as progenitors for neuroendocrine cells, ciliated epithelial cells, and Club cells in the bronchi/bronchioles, while the distal epithelial cells, defined by expression of Sox9, Id2, and Sftpc in fetal lungs, are the progenitors of peripheral alveolar epithelial cells ^24^. The mechanisms by which mesenchymal cells regulate airway epithelial cell fate during lung development are largely unknown. Interestingly, in the *Bmpr1a* CKO lung, Sox2-positive epithelial cells were barely detected in the proximal airways. Similarly, lack of further differentiation of Foxj1-positive cells was also observed. In contrast, cells expressing Sox9 and Sftpc, which are normally confined to the distal epithelial cells during lung branching, were expanded to the proximal region of the airways in *Bmpr1a* CKO lungs. The analysis of tissue RNA-Seq data revealed that the mesenchymal *Bmpr1a* knockout downregulated multiple proximal airway epithelial marker genes, while various distal airway epithelial marker genes were upregulated (Fig. 7B). Ectopic expression of Bmp4 in fetal mouse lung has been reported to cause distal epithelial expansion ^15^. Increased expression of *Bmp4* in E15.5 *Bmpr1a* CKO lungs was initially found by RNA-seq (LogFC>1, GEO#: GSE97946) and validated by RT-PCR (Fig.7C). The inverse relationship between Bmpr1a deletion (intracellular BMP signaling) and Bmp4 ligand expression in these fetal lung mesenchymal progenitor cells (MPCs) was further confirmed by comparing Bmp4 expression in isolated fetal lung mesenchymal cells with different *Bmpr1a* genotypes (Fig. 7D-7E). Alterations of other key pathways including Fgf, Wnt, and Shh were not detected by RNA-Seq and RT-PCR through analyzing the related targeted genes.

**Fig. 7.**
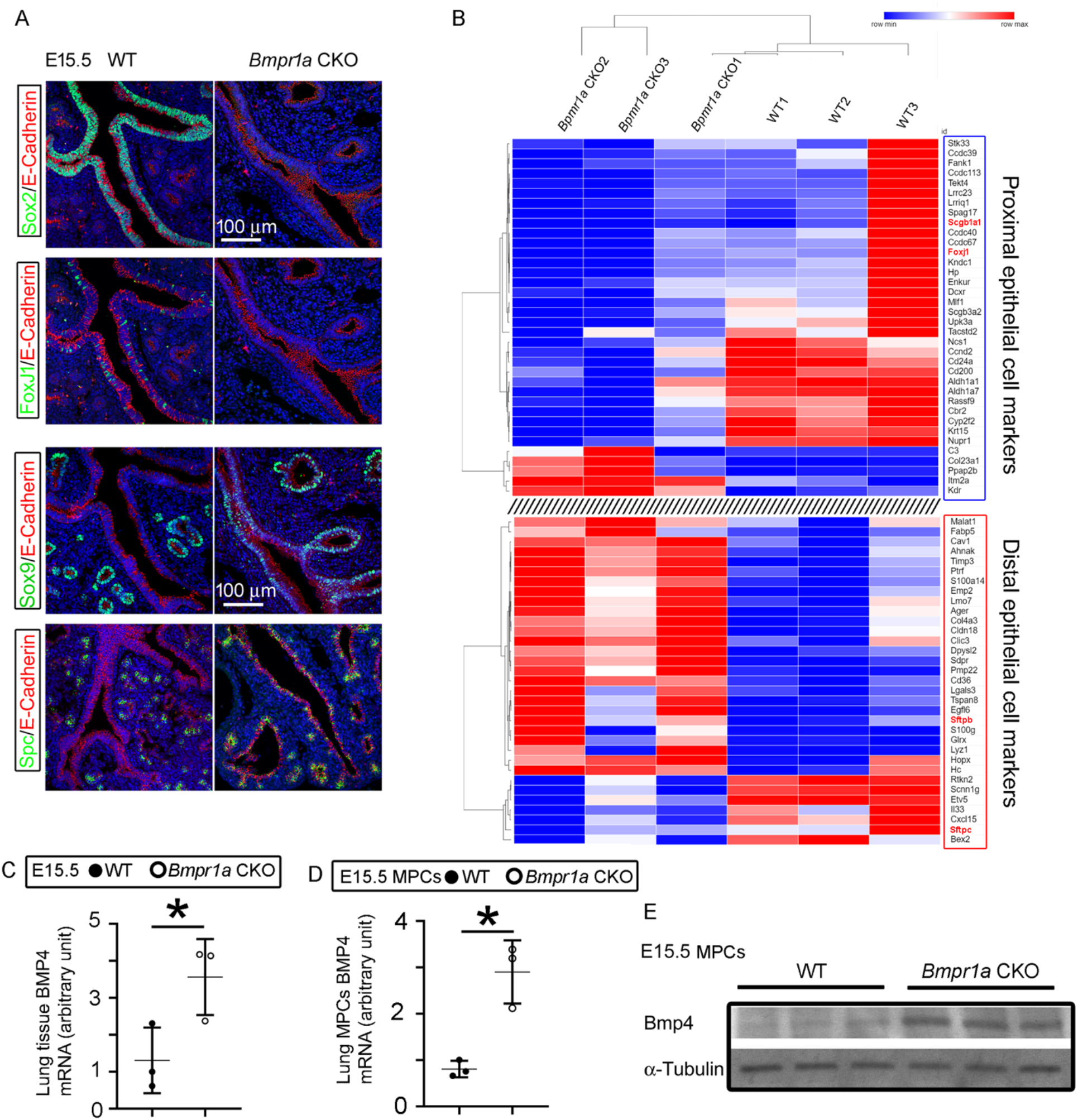
Mesenchymal *Bmpr1a* deletion disrupted airway epithelial proximal-distal differentiation and development. (A) Proximal epithelial cells, marked by Sox2 and Foxj1 staining, were significantly decreased in the proximal portion of the airways in E15.5 *Bmpr1a* CKO lungs. Ectopic distribution of distal epithelial cells marked by Sox9 and Spc staining was detected in the proximal airways of E15.5 *Bmpr1a* CKO lungs. (B) Heatmap of RNA-Seq data showing significant changes in the marker genes of proximal and distal epithelial cells. (C) Increased *Bmp4* expression at the mRNA level was detected in E15.5 *Bmpr1a* CKO lung tissue by RT PCR. (D & E) Bmp4 expression in isolated fetal lung mesenchymal cells with genotypes of WT vs. *Bmpr1a* CKO was analyzed at both the mRNA and protein levels by RT PCR and WB respectively.

## Discussion

Our current study provides solid in vivo evidence for the first time that Bmpr1a-mediated BMP signaling in lung mesenchymal cells is indispensable in lung development. Lung mesenchymal deletion of *Bmpr1a* at an early developmental stage causes abnormal branching morphogenesis and then severe lung cysts at late gestation in mice. Interestingly, congenital pulmonary cysts are reported in human congenital pulmonary airway malformation (CPAM) patients as a result of abnormal prenatal lung development ^25^. Reduction of airway SMCs and decreased subepithelial elastin fibers are also found to be the common pathological changes in the cystic airways of human CPAM samples ^17^. Although the congenital pulmonary cysts in CPAM patients can be identified by a prenatal ultrasound examination, the tissue specimens for pathologic analysis are only available from postnatal surgery, when advanced airway cysts have already developed. This makes it difficult to study the pathogenic molecular mechanisms that trigger early cyst formation during fetal lung development in human subjects. Studies in mouse models suggest that altered growth factor signaling in a prenatal time window is sufficient to cause cyst formation ^26^. The abnormal BMP signaling pathway may be involved in the initiation of CPAM, but not in the later stage when cystic lesions have formed. Interestingly, integrated suppression of BMP signaling pathway was recently found in human CPAM specimens by an RNA-seq approach ^27^. Therefore, the lung mesenchymal *Bmpr1a* knockout model developed here is a potentially useful tool for studying the pathogenic process of congenital pulmonary airway malformation.

It has been reported that epithelial overexpression of Bmp4 appears to induce extensive mesenchymal SMC differentiation in vivo ^28^, and that BMP4 promotes myocyte differentiation and inhibits cell proliferation in cultured human fetal lung fibroblasts via the Smad1 pathway ^29^. Smad1 and Smad5 are critical downstream effectors of BMP type I receptors, mediating BMP canonical signal pathway ^9,30^. However, our in vivo study indicates that simultaneous deletion of *Smad1* and *Smad5* in mouse lung mesenchyme does not affect the differentiation of airway SMCs although it disrupts early embryonic lung morphogenesis. This suggests that the decrease in phosphorylated Smad1 and Smad5 in the *Bmpr1a* CKO lungs is not responsible for the defective airway SMC growth. In addition to the Smad-dependent BMP signal pathway, BMP-activated Bmp receptors also function through mitogen-activated protein kinases (MAPK) ^9,30,31^. Many studies have shown that p38 MAPK regulates cell proliferation in a variety of smooth muscle cell types, including airway SMCs ^32,33^. The p38 MAPK is also involved in myogenic differentiation and myofibroblast trans-differentiation ^34-36^. In the present study, reduced phosphorylation of p38 was detected in mesenchymal *Bmpr1a* CKO lungs while addition of BMP4 activated the p38 signal pathway in mouse primary fetal lung mesenchymal cells. Moreover, the myogenic response to BMP4 stimulation in the cultured fetal lung mesenchymal cells could be blocked by treating the cells with a p38 inhibitor. This suggests that p38 is the primary intracellular signaling component to mediate BMP’s regulatory effects on airway myogenesis.

Despite being commonly characterized by the expression of a variety of myogenesis-associated genes, airway and vascular SMCs seem to originate from distinct mesenchymal progenitor populations^16^, and are regulated by distinct molecular mechanisms. Our study reveals that *Myocd* is a prominent target gene that links Bmpr1a to the genes associated with airway SMC development. However, Bmpr1a, p38 and Myocd are not critical in fetal pulmonary vascular SMC development, and inhibition of these molecules in vivo and in vitro does not have any observable effect on vascular SMCs in fetal lungs. Our previous study has demonstrated that *Myocd* is abundant in airway SMCs but barely expressed in vascular SMCs ^18^, and that cardiovascular smooth muscle cell development is mediated by myocardin related transcription factors, the other members of *Myocd* family ^22^. Distinct regulation between airway and vascular SMCs is also evidenced by the phenotypes of mice with various mutations ^28,37^. For instance, hypomorphic FGF-10 mice only display decreased expression of airway smooth muscle actin ^38^, while deletion of Wnt-7b selectively disrupts the development of vascular SMCs ^39^. Although reduced expression of Bmpr1a is associated with various forms of acquired as well as primary nonfamilial pulmonary hypertension ^40^, the role of Bmpr1a-mediated signaling in regulating prenatal lung vascular SMC development has not been studied. Our current data suggest that Bmpr1a is dispensable in fetal vascular SMC development.

Previous studies have suggested that airway smooth muscle cells initially develop from local mesenchymal cells around the tips of epithelial buds undergoing bifurcation. These cells wrap around the bifurcating cleft and neck of the terminal buds, and then grow rigidly alongside the elongating bronchial tree ^20,28,41^. Inhibition of SMC differentiation in embryonic lung explant culture, such as by blocking L-type Ca^2+^ channels using nifedipine, or by inhibiting FGF and SHH signaling using SU5402SHH and cyclopamine, prevented airway terminal bifurcation ^20^. However, our current and a recently published study demonstrate that abrogation of airway SMCs by deleting mesenchymal *Myocd* does not cause any significant abnormalities in airway branching morphogenesis. This suggests that loss of SMCs is neither the primary nor sole mechanism accounting for the reduction of airway branching, dilation of terminal airways, and development of airway cysts in our *Bmpr1a* CKO lungs. Notably, the airway subepithelial elastin defects were uniquely found in the *Bmpr1a* knockout lungs with extensive cystic lesions but not in the *Myocd* CKO lungs. Interestingly, elastin expression in the developing lung parenchyma prior to alveologenesis is predominantly localized to the mesenchyme in the developing airways, as indicated by our elastin immunostaining data, suggesting that elastin plays a role in airway branching. Previous study has revealed that the perinatal development of terminal airway branches is arrested in mice lacking elastin (*Eln* -/-), resulting in dilated distal air sacs that form abnormally large cavities ^42^. It is worthy to note that airway smooth muscles are normal in these elastin null mice, implying that loss of elastic fiber alone is not sufficient to cause airway cystic pathology, but could serve as an additional factor contributing to the lesions in vivo. This assumption is also supported by the fact that congenital airway cysts in the lungs of human CPAM patients are characterized by compromised smooth muscle and elastin fiber^17^. Future studies are needed to investigate whether the combined defects of airway smooth muscle cells and elastin fibers are the pathogenic mechanisms behind congenital airway cyst formation.

The upregulation of multiple Bmp ligands, particularly Bmp4, in the *Bmpr1a* CKO lung tissues as compared with the WT lungs is a complex scenario. The isolated fetal lung mesenchymal cells of *Bmpr1a* CKO mice also showed increased Bmp4 expression as compared to the WT lung cells. This might be due to an autoregulatory feedback loop between Bmp ligand expression and intracellular BMP signaling. Previous studies have shown that disruption of BMP signaling by LDN-193189 can substantially improve Bmp4 production ^43^, and that deletion of Bmpr1a and Acvr1 in the lens-forming ectoderm increases Bmp2, 4 and 7 transcripts ^44^. These in vitro and in vivo studies clearly suggest the inverse relationship between mesenchymal BMP signaling activity and Bmp ligand expression. Misexpression of BMP4 results in a decrease of Sox2^+^ proximal progenitors and an expansion of Sox9^+^/Id2^+^ distal progenitors in fetal mouse lungs, leading to malformed airways ^45^. Consistently, overexpression of Bmp antagonist Xnoggin or gremlin in fetal distal lung epithelium leads to a severe reduction of distal epithelial cell types and a concurrent expanded expression of the proximal epithelial cell markers CC10 and Foxj1 ^14,46^. In our *Bmpr1a* CKO lungs, the proximal-distal patterning of lung airways is also substantially disturbed, as indicated by the aberrant distribution of proximal and distal epithelial cells. Evidence increasingly suggests that the lung mesenchyme is closely involved in respiratory lineage specification and epithelial differentiation by producing many critical signaling molecules, such as Bmp ligands, that direct endodermal expression of specific cell markers in a temporal and spatial fashion ^1,2^. Therefore, the decrease in proximal cell types and ectopic expression of distal cell markers (Sox9) in the *Bmpr1a* CKO lungs may be subsequently caused by abnormal increase of Bmp ligands. This aberrant epithelial growth may also contribute to airway cyst formation, as multiple mouse models have suggested that loss of Sox2^+^ or expansion of Sox9^+^ in airway epithelia are associated with altered epithelial differentiation and branching, as well as cystic pathology in fetal lungs ^45,47,48^.

In summary, mesenchymal Bmpr1a is essential for lung development, particularly airway branching morphogenesis. Disruption of Bmpr1a-mediated mesenchymal signaling causes prenatal airway malformation and cystic lesions. Multiple mechanisms, including deficiency of airway smooth muscle, airway elastin fiber defect, and perturbation of the Sox2-Sox9 epithelial progenitor axis, may contribute to the congenital airway cystic pathogenesis. In addition, the downstream Smad-independent p38 pathway appears to be critical in mediating BMP-regulated airway smooth muscle development. Our study helps to understand both normal lung development and the mechanisms of related airway malformation and congenital pulmonary diseases.

## Materials and Methods

### Mice

The *Tbx4-rtTA*/*TetO-Cre* transgenic line was generated in our lab ^16^. *Floxed*-*Bmpr1a* (*Bmpr1a ^fx/fx^*) mice were provided by Dr. Yuji Mishina ^49^. *Floxed*-*Myocd* (*Myocd ^fx/fx^*) mice were provided by Dr. Michael S. Parmacek ^50,51^. *Floxed*-*Smad1* (*Smad1^fx/fx^*) mice were originally generated by Dr. Anita Roberts ^52^. *Floxed*-*Smad5* (*Smad5 ^fx/fx^*) mice were provided by An Zwijsen at KU Leuven ^53^. *Cspg4*- *DsRed* mice were purchased from The Jackson Laboratory (#008241). *Tagln-YFP* reporter mice were provided by Dr. Jeffrey Whitsett at Cincinnati Children’s Hospital. To generate the lung mesenchyme-specific conditional knockout (CKO) mice of genes studied in this work, *Bmpr1a^fx/fx^*, *Myocd ^fx/fx^*, and *Smad1^fx/fx^/Samd5^fx/fx^* mice were bred to the *Tbx4-rtTA/TetO-Cre* mice carrying *Bmpr1a^fx/+^*, *Myocd ^fx/+^* or *Smad1^fx/+^/Samd5^fx/+^* respectively. Doxycycline (625 mg/kg in food (TestDiet) and 0.5 mg/ml in drinking water (Sigma)) was given to the mothers from E6.5 to induce Cre-mediated floxed gene deletion. The littermate controls were the mice without any floxed-gene deletion due to lack of transgenes *Tbx4-rtTA* and/or *TetO-Cre*. *Tagln-YFP* mice were crossed with *Cspg4-DsRed* mice to generate double fluorescence reporter mice. Timed mating was set up as previously described ^12^. Mouse fetal lungs of different embryonic stages were collected from pregnant females. All mice were bred in C57BL/6 strain background. The mice used in this study were housed in pathogen-free facilities. All mouse experiments were conducted in accordance with NIH Guide and approved by the Institutional Animal Care and Use Committee of Children’s Hospital Los Angeles.

### Cell isolation, culture, and treatment

Feal lung mesenchymal cells from E15.5 lungs were isolated and cultured as previously described^54^. To study the role of BMP signaling in subpopulations of SMCs, lung mesenchymal cells from *Tagln-YFP*/*Cspg4-DsRed* reporter mice were further sorted into perivascular SMCs and non-perivascular SMCs based on their fluorescent reporter expression using FACS (BD, FACSAria I). The cells were then plated at low density (∼10^4^/100-mm dish) and cultured in αMEM medium supplemented with 20% FBS, 2 mM L-glutamine, 55 μM 2-mercaptoethanol and antibiotics (100 U/ml penicillin and 100 μg/ml streptomycin). The cells (<10 passages) were treated with recombinant mouse BMP4 protein (50 ng/ml), LDN193189 (200 nM) or p38 pathway inhibitor (SB203580, 1 µM) for two weeks. BMP4 protein was purchased from R&D Systems (5020-BP). LDN193189 was provided by Dr. Paul Yu at Harvard Medical School ^55^, and SB203580 was purchased from Sigma (#S8307).

### Histology and immunofluorescence analysis

Fetal mouse lungs were isolated and imaged under a dissecting microscope. Branching tips were quantified manually. For the size of branching tips, a freeform trace around each airway tip was drawn and the area within the trace was measured using ImageJ software. The data were presented as relative changes of Bmpr1a CKO to the littermate controls ^56^. For preparation of regular paraffin tissue sections and 2-D histological examination, isolated fetal lungs were fixed in 4% paraformaldehyde, as described in our previous publication ^12^. Hematoxylin and eosin (H&E) and immunofluorescent staining was performed as described previously ^5^. For whole mount immunofluorescence staining and 3-D imaging, fetal lungs were isolated and fixed in DMSO:methanol (1:4) overnight at 4 °C, following a protocol described before ^8^. Antibodies used for immunofluorescent staining were listed in the Suppl. Table 1.

### Cell proliferation and apoptosis analyses

EdU incorporation was used to assess the in vivo cell proliferation following instruction from the manufacture (#C10337, Thermo Fisher Scientific). Cell quantification was performed using Fiji imaging software (1.52p) as described previously ^5^. Apoptosis was evaluated by TUNEL assay (#S7110, MilliporeSigma). E15.5 lung specimens with additional treatment of DNase I served as the positive control.

### RNA Extraction and RNA-Seq analysis

E15.5 mouse lungs were dissected in RNase-free PBS and flash-frozen in liquid nitrogen. After genotyping, mRNA from WT and *Bmpr1a* CKO lungs was prepared by using the TRIzol RNA extraction reagent (ThermoFisher, #15596026). The RNeasy Micro Kit (Qiagen, #74004) was then used to clean up the extracted mRNA. Total RNA quality was measured with a bioanalyzer (Agilent) with RIN greater than 7.0 being acceptable. RNA-sequencing libraries were generated and sequenced by Quickbiology (Quickbiology Inc. Pasadena, CA), using the Illumina TruSeq v2 kit (Illumina, San Diego CA). Data processing was performed using the USC high performance-computing cluster (https://hpcc.usc.edu/). Roughly 50 million 100bp single-end sequences were aligned to the Gencode M8 annotation ^57^ based on the Genome Reference Consortium mouse genome (GRCm38.p4) and using the STAR aligner ^58^. Read counts per gene were calculated using the HTSeq-count software ^59^. Differential gene expression was determined using the R/Bioconductor software ‘edgeR’ ^60^. Genes were considered significantly different if their false discovery rate (FDR) corrected P-values were less than 0.05 and Log2-Fold change was greater than 1.0. The sequencing data was deposited to the NCBI GEO repository with accession number GSE97946.

### Real-time PCR and western Blot

Lung mesenchymal cells and tissue specimens were harvested and flash-frozen in liquid nitrogen for the subsequent mRNA and protein analysis. Total RNA was isolated from the cultured cells and lung tissues using the RNeasy Kit (Qiagen, #74106). The cDNA synthesis and real-time PCR were performed as described in a previous publication ^5^. The related oligonucleotide primers are listed in Suppl. Table 2. Protein lysate was prepared from the cells and lung tissues (n>3 per group) and analyzed by western blot as previously described ^54^. The related antibodies used for western blot are listed in Suppl. Table 1.

### Statistical Analysis

The quantitative data are presented as means ± SD. All in vitro experiments were repeated at least three times, and data represent consistent results. At least 5 mice per experimental group were utilized and the statistical difference between any two groups was assessed by an independent-sample t-test. One-way analysis of variance (ANOVA) was used to examine differences between groups of three or more, followed by *post-hoc* comparisons to study differences among individual group. Chi-square test with Yates correction was used for GO enrichment analysis. *P*< 0.05 was considered statistically significant.

## Acknowledgements

We thank Dr. Robert Mecham at Washington University St. Louis, Dr. Paul Yu at Harvard University, Dr. Jeffrey Whitsett at Cincinnati Children’s Hospital, and Dr. An Zwijsen at KU Leuven for providing the mouse lines and other reagents. Dr. Esteban Fernandez at the Cell Imaging Core of Children’s Hospital Los Angeles helped with the confocal imaging. This study was supported by NIH grants HL151699 and HL141352 (W.S.).

## Notes

### Competing Interest Statement

The authors have declared no competing interest.

### Summary of Updates

Figure 1, 2, 6, and 7 revised with addition of new data. The result and discussion sections updated to clarify the quantitative changes of lung cysts, airway smooth muscle cell defect, and the relationship between Myocd deletion and Bmpr1a expression. Supplemental Figure 1 revised.

