## Supplemental figures and tables for "Defective mesenchymal Bmpr1a-mediated BMP signaling causes congenital pulmonary cysts"

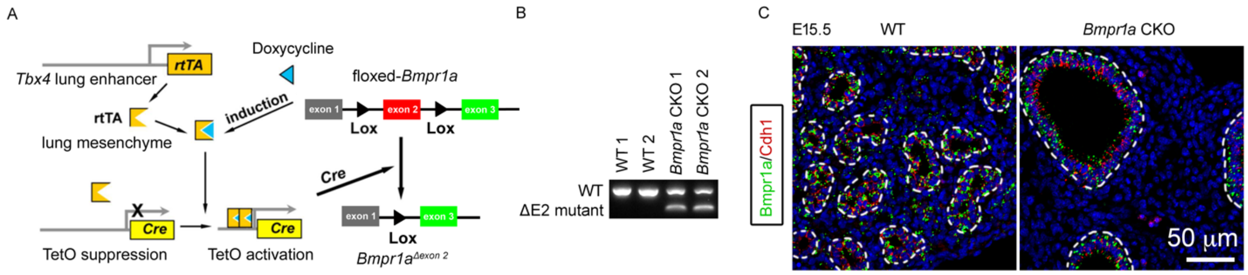


**Supplementary Fig. 1.** *Bmpr1a* was specifically deleted in lung mesenchymal cells. A. Schematic representation of the lung mesenchyme-specific knockout of *Bmpr1a* by *Tbx4-rtTA*/*Teto-Cre* driver line. B. Verification of Bmpr1a genetic deletion at the mRNA level. △E2: exon 2 deletion. (C) Immunostaining of Bmpr1a (green) and Cdh1 (red). The airways are highlighted with dot lines.

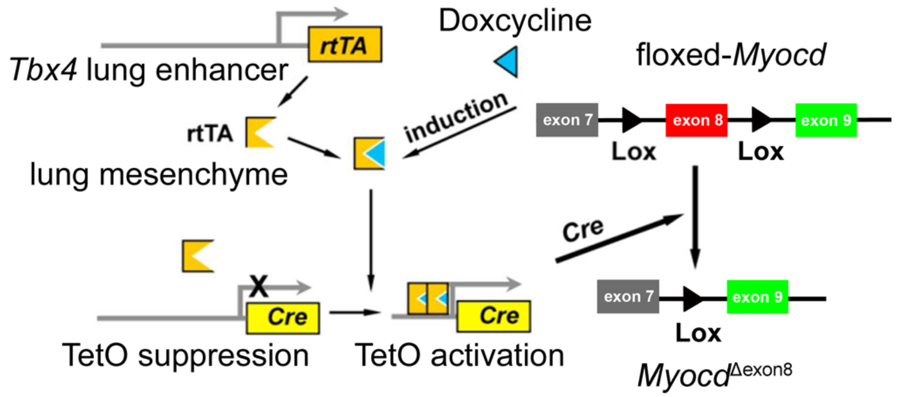


**Supplementary Fig. 2.** Schematic representation of the lung mesenchyme-specific knockout of *Myocd* by *Tbx4-rtTA/Teto-Cre* driver line.

**Supplementary Table 1:** Primary antibodies used for immunochemistry & western blot

| Antibody name | Vendor | Catolog # |
|---|---|---|
| Goat anti-Bmpr1a | Santa Cruz, | sc-5676 |
| Mouse anti-Cytokeratin | Sigma | C2562 |
| Rat anti-E-cadherin | ThermoFisher Scientific | 13-1900 |
| Mouse anti-SMA | Sigma | A2547 |
| Rabbit anti-Elastin | Generated by Dr. Robert Mecham at Washington University, St. Louis (Luo et al., 2018; Young et al., 2020) | |
| Goat anti-Pecam | Santa Cruz | Sc-1506 |
| Rabbit anti-NG2 chondroitin sulfate proteoglycan | Millipore | AB5320 |
| Rabbit anti-Laminin | ThermoFisher Scientific | RB-082 |
| Goat anti-Collagen III alpha 1 | Novus Biologicals | NBP1-26547 |
| Mouse anti-Foxj1 | | |
| Rabbit anti-Sox2 | Seven Hills Bioregeants | WRAB-1236 |
| Rabbit anti-Sox9 | Cell Signaling Technology | 82630 |
| Rabbit anti-Sftpc | Seven Hills Bioregeants | WRAB-9337 |
| T1α | Developmental Studies Hybridoma Bank | 8.1.1 |
| Rabbit anti-Myh11 | Millipore | MABT464 |
| Rabbit anti-phospho-Smad1/5 | Cell Signaling Technology | 9516 |
| Rabbit anti-Smad1 | Cell Signaling Technology | 6944 |
| Rabbit anti-phospho-p38 | Cell Signaling Technology | 4511 |
| Rabbit anti-p38 | Cell Signaling Technology | 9212 |
| Rabbit anti-phospho-Erk1/2 | Cell Signaling Technology | 4370 |
| Rabbit anti-Erk1/2 | Cell Signaling Technology | 4695 |
| Rabbit anti-phospho-Jnk | Cell Signaling Technology | 4668 |
| Rabbit anti-Jnk | Cell Signaling Technology | 9252 |
| Mouse anti-GAPDH | Fitzgerald | 10R-G109a |

**Supplementary Table 2:** Primers used for real-time RT-PCR

| Gene | Oligonucleotide DNA sequence |
|---|---|
| *Bmpr1a*-△exon2 | 5'- GGG AGC CTG TCT GTT CAT CA -3' |
| | 5'- TTT CGG TGA ATC CTT GCA TT -3' |
| *Acta2* | 5'- AAT GCA GAA GGA GAT CAC GG -3' |
| | 5'- TCC TGT TTG CTG ATC CAC ATC -3' |
| *Myh11* | 5'- AGA AGG AGC GAA ACA CAG AC -3' |
| | 5'- TGT CAC ATT AAT CCC CAC GAG -3' |
| *Cnn1* | 5'- TGG CAC CAG CTG GAG AAC AT -3' |
| | 5'- TCA AAC AGG TCG TTG GCC TCA -3' |
| *Tagln1* | 5'- CCA GAC TGT TGA CCT CTA TGA AG -3' |
| | 5'- TCT TAT GCT CCT GGG CTT TC -3' |
| *Cspg4* | 5'- CCT TCA CGA TCA CCA TCC TTC -3' |
| | 5'- AAT CAT TGT CTG TTC CCC TGA G-3' |
| *PECAM1* | 5'- GAG ATG TCC AGG CCA GCT G -3' |
| | 5'- CTC ACT GTA CAC CGT CTC TG -3' |
| *Lama1* | 5'- AAA GGA AAG TGT CAG TAC CAG G-3' |
| | 5'-TTC TCT AAG CAT CGC AAG GG-3' |
| *Lama2* | 5'-GTC TGG GAT CAT TCT CTT GGG-3' |
| | 5'-TTT CCT CAT TGT CCG TGT CC-3' |
| *Col3a1* | 5'-GAA GTC TCT GAA GCT GAT GGG-3' |
| | 5'-TTG CCT TGC GTG TTT GAT ATT C-3' |
| *Eln* | 5'- ACT TTC TCC CAT TTA TCC AGG TG -3' |
| | 5'- AAG ATC ACT TTC TCT TCC GGC -3' |
| *Foxj1* | 5'- CCA CCT GGC AGA ATT CCA T -3' |
| | 5'- CCT CCG CTT CTT GAA GGC -3' |
| *Sox2* | 5'- TTT GTC CGA GAC CGA GAA GC -3' |
| | 5'- CTC CGG GAA GCG TGT ACT TA-3' |
| *Sox9* | 5'- CAA GAC TCT GGG CAA GCT-3' |
| | 5'-GGG CTG GTA CTT GTA ATC GG-3' |
| *Spc* | 5'- CTT TCG CTA GAA AAC TCC AGA ACT -3' |

| | |
|---|---|
| | 5'- AAG AAT CGG ACT CGG AAC CAG -3' |
| *Myocd* | 5'- CGA TCA GTC TTA CAG TTA CGG C -3' |
| | 5'- CTC AGG GAA TCT TCA GTC TTG G-3' |
| *Gapdh* | 5'-GGT GGA GCC AAA AGG GTC AT-3' |
| | 5'-AGT TGT CAT ATT TCT CGT GGT TCA-3' |
